# A high-resolution cryo-EM structure of a bacterial M-protein reveals a compact structure that diverges from related M-proteins

**DOI:** 10.1101/2023.09.18.558297

**Authors:** Bradley M. Readnour, Sheiny Tjia-Fleck, Nathan R. McCann, Yetunde A. Ayinuola, Francis J. Castellino

## Abstract

The surface of *Streptococcus pyogenes* (GAS) is studded with virulence determinants, with the most abundant being the characteristic M-protein used to serotype various strains of the bacterium. There are >250 strains of GAS serotypically distinguished by their M-proteins. Major pathogenic mechanisms of GAS require that this microorganism hijacks host components for survival, many of which are involved in hemostasis. One of these processes involves the binding of human host plasminogen (hPg) to an abundant GAS M-protein receptor (PAM). When bound to PAM, hPg is readily activated to the serine protease plasmin (hPm) by bacterial and host hPg activators, and cell-bound hPm is protected from inactivation by its natural inhibitors. This stabilizes a potent protease on GAS cells which aids in their survival and dissemination. Highly evolutionary domain-related M-proteins are assumed to form long alpha-helical projections, without tertiary structure, although no M-protein complete structure has been determined. Here, we employed cryogenic electron microscopy to solve such a structure anchored to a lentivirus particle membrane. Contrary to the belief in this field that M-proteins are extended long tropomyosin-like coils, we show that PAM folds through intra- and inter-domain interactions to a much more globular form on the cell surface. The nature of the folding and the many interactions involved in forming the PAM tertiary structure are summarized herein.

**Significance:** We provide a unique approach to solve high-resolution structures of *Streptococcus pyogenes* (GAS) M-proteins, abundant virulence determinants on the GAS surface. Because of their unusual nature, no full high-resolution structure of any M-protein has been determined, especially when membrane-bound. Herein, we provide a unique general methodology for solving these structures by engineering a M-protein to be anchored to a lentivirus particle membrane for effective use in cryo-EM. Using this approach, we provide the first structure of a complete bacterial M-protein and show, that this M-protein is a monomeric globular structure on the cell surface, and not a dimeric coiled-coil, as generally believed. Thus, individual M-proteins may adopt structures that have evolved to accommodate their major host binding partner.

## Introduction

Plasminogen receptors (PgRs) are present on the surface of a variety of cell types and play an important role in the lysis of blood clots within the human body and in surveillance of fibrin *via* functional binding of the zymogen, plasminogen (hPg) and its activated product, plasmin (hPm) (1, 2). However, the ability to bind hPg and hPm can also be used pathologically by invasive cells to allow them to disseminate and invade deep tissue sites (3). One of the most prominent cases of bacterial cells using hPg to aid in the invasion of host cells occurs in group A *streptococcus pyogenes* (GAS). In some specialized GAS strains, M-protein (M-Prt), the most abundant virulence factor on the surface of GAS, has the ability to directly bind hPg to the cell surface (4). Bound hPg can then be readily activated by streptokinase (SK), a secreted endogenous hPg activator (5), or even host hPg activators. As a result, bound hPg is converted to the active serine protease hPm on the cell surface, while, unlike soluble hPm, is resistant to naturally occurring inactivators and inhibitors (6). By disrupting innate immune barriers, *e.g.,* tight cellular junctions, the spread of hPm-adorned GAS cells throughout the body occurs with a greatly increased rate of infection.

Of the approximately 250 strains of GAS that are serotypically unique, M-Prts are a dominant virulence determinant. A subgroup of GAS cells contains a type of M-Prt, hPg binding group A streptococcal M-Prt (PAM), of which there are a number of naturally occurring sequence variants, all of which bind directly to hPg with high affinity (6). Other subgroups of M-Prts have been shown to bind to fibrinogen directly (7) which then recruits hPg to the surface where it is activated to hPm. PAM-type M-Prts have been shown to have the highest binding affinity for hPg (K_d_ ∼1 nm) (8) when compared to other putative receptors within GAS cells, *viz.,* α-enolase (SEN) or GAPDH (9, 10). This high binding affinity, natural protection, and high surface abundance makes PAM-type M-Prts an important target for research in understanding the interactions between hPg and GAS cells that result in its pathogenicity.

Despite the significant roles that M-Prts play in the virulence of GAS cells, little is known about their three-dimensional structures. The large number of flexible regions within M-Prts limit their ability to crystalize, and its molecular weight (∼41 kDa) makes it a difficult target for high resolution structure-based NMR (too large) and cryo-EM (too small). Fragments of PAM from GAS strain AP53 (PAM_AP53_) have been modeled using NMR (11, 12) and X-ray crystallography (13), but to date, the full structure of any type of M-Prt has never been solved. In the case of PAM, a complete structure is necessary in order to understand how this M-Prt interacts with hPg and the role it plays in GAS virulence.

In this communication we uniquely designed a method of integrating a bacterial surface protein into the membrane of a lentiviral particle (LVP) to allow for cryo-EM imaging of low molecular weight membrane proteins. As a result, we have obtained a 3.8 Å high-resolution map of PAM_AP53_ actively bound to hPg on the cell surface. This map was used to design a *de novo* model of PAM bound to hPg on the cell surface. This structure differs significantly from any previously proposed structures or AI-generated models of M-proteins using programs such as AlphaFold (14, 15). This structure will play an important role in understanding the interactions between PAM and hPg and the role of PAM in the overall surface structure of GAS cells. The method of using LVPs in the imaging of low molecular weight bacterial membrane proteins will play an important role in further design of protein structures in the future.

## Results

### Surface expression of PAM on lentiviral particles (LVP)

Expression of the PAM construct **(Fig. 1)** on the surface of a LVP allows for a membrane anchor to increase the contrast of the protein for cryo-EM imaging, as well as to obtain the structure of PAM in a more natural state than PAM in solution. To enable the surface expression of PAM, we exchanged the signal sequence (SS) and transmembrane + cytosolic regions (TMD+Cy) of PAM with those of a glycoprotein (VSV-G) of vesicular stomatitis virus.

**Figure 1.**
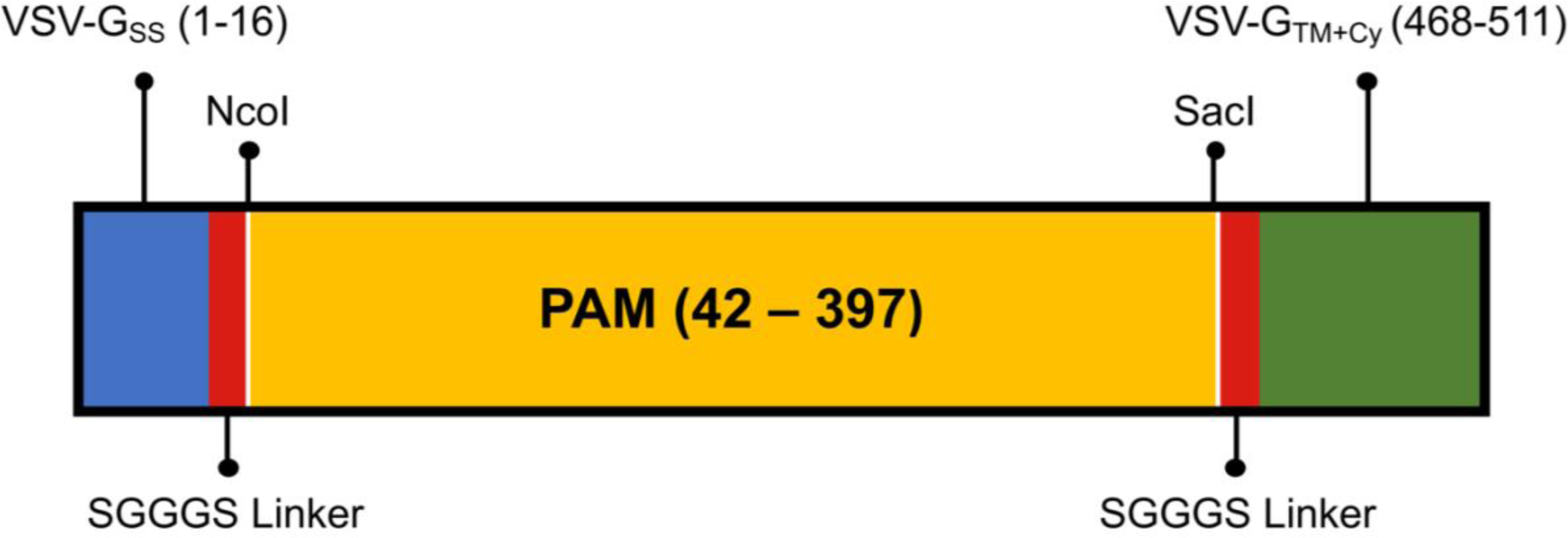
Construction of the expression vector for PAM_AP53_ on LVP particles. A synthetic expression plasmid was generated containing the codons for the 16 amino acid residue 5’-signal sequence (SS) of vesicular stomatitis virus-G (VSV-G) followed sequentially by codons for a short linker sequence, a NcoI restriction site, a SacI restriction site, another linker sequence, and the 3’- transmembrane (TM) domain + cytosolic tail (Cy) of VSV-G_TM+Cy._ The mature open reading frame of PAM, encoding the cDNA for residues 42-397 (1-356 of the mature protein without the 41 amino acid residue SS) and COOH-terminal TM+Cy tail of PAM, also containing 5’ Nco1 and 3’ SacI restriction sites, was cloned from the PAM gene and inserted into the expression plasmid *via* the NcoI and SacI restriction sites. The diagram illustrates the final PAM_AP53_ expression vector that was transduced into the LVP allowing for stable expression of PAM_AP53_ on the LVP membrane.

As shown in **Figure 2**, our PAM-LVP preparations gave a positive signal with anti-PAM generated in-house against the PAM amino-terminal region, thus confirming the surface expression of PAM on the LVP. This signal intensity is comparable to the signal obtained for 50 ng of free PAM (recombinant PAM in solution) indicating that the PAM level on the LVP surface is similar to that determined previously to be ideal for cryo-EM imaging of this protein. The SARS-CoV-2 Spike Protein-LVP negative control (a component of the kit) gave no signal. Hence, the signal obtained from PAM-LVP was not due to the non-specific interaction of the viral proteins with anti-PAM.

**Figure 2.**
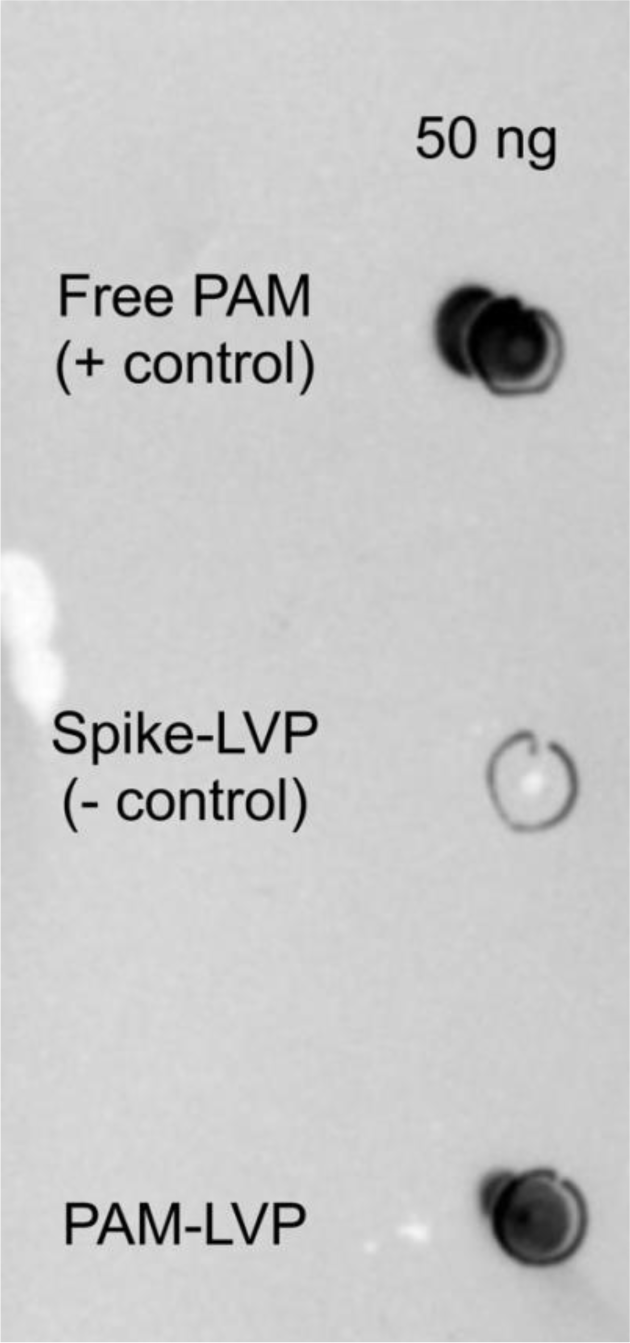
Dot blot of PAM-LVP. To confirm the existence of PAM_AP53_ on the surface of the LVP (PAM-LVP) a dot blot was performed. Both free PAM_AP53_ and PAM-LV were added to a dot blot membrane, washed, and incubated with an in-house anti-PAM generated against the N-terminal region of PAM_AP53_ (HVR-A-domain-partial B-domain). PAM_AP53_ was used as the positive control and applied at the same concentration that was used in cryo-EM imaging. This confirmed that PAM was on the surface of the LVP at a similar concentration to the free PAM used in cryo-EM. LVP with SARS-Cov2 Spike Protein (Spike-LVP) was used as a negative control.

### High-resolution structure of PAM bound to hPg on the surface of LVP

Micrographs containing PAM-LVP were taken using the Titan Krios. The PAM-LVPs show some variation in size, between 80-120 nm, with the majority being unilammelar ∼100 nm vesicles (**Fig. 3A**). Moreover, the PAM concentrations on each LVP also vary, with some viruses containing little to no PAM (**Fig. 3B**) and others having abundant PAM_AP53_ protein (**Fig. 3C**). The circular structure of the LVP allows for multiple orientations of PAM_AP53_ on the surface of the LVP to be imaged (**Fig. 3D**).

**Figure 3.**
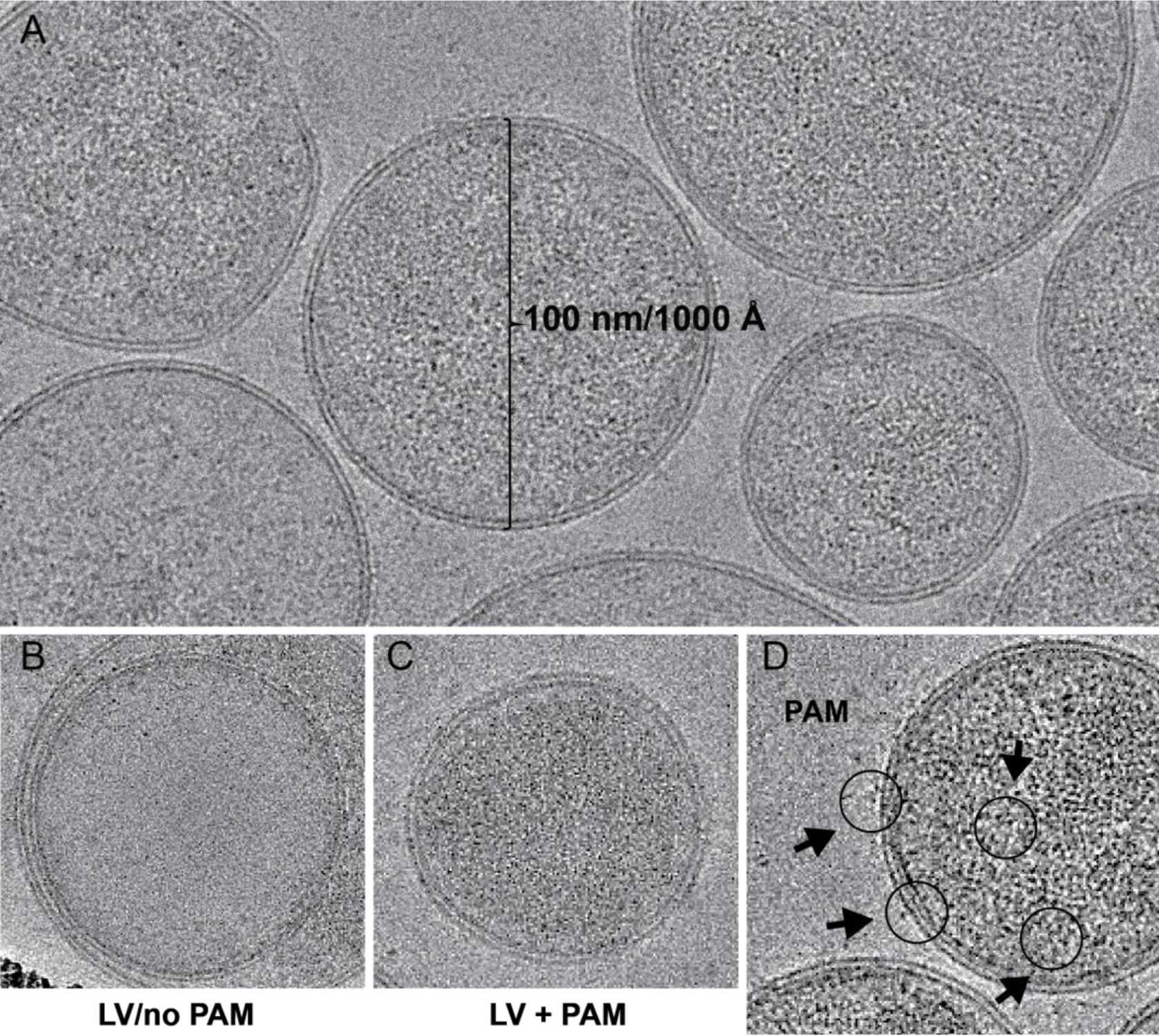
Micrograph of LVPs with PAM_AP53_ bound to the surface. **(A).** Cryo-EM imaging confirms the existence of largely uniform 100 nm LVs. **(B).** Empty LVP micrographs were removed manually. **(C).** In comparison with **B**, the LVP with large amounts of PAM on the surface are clearly visible. **(D.)** Examples of PAM on the LVP are marked with circles and arrows. These micrographs were the basis of the final cryo-EM structure. Proteins on the surface were isolated by Cryosparc and were used for the 3D map.

A total of 6,544 micrographs were imaged over the course of two 24 hr runs. A final total of 4,054 micrographs were used for final map reconstruction with the remaining micrographs removed due to the presence of crystalline ice, or a lack of LVP. A low specificity particle selection was performed searching for particles with a minimum radius of 50 Å and a maximum radius of 100 Å, with a box size of 320 pixels and pixel size of 0.437 Å. This resulted in an initial particle set of ∼1,200,000 particles. These particles were used for an initial 2D classification containing 50 classes. From these classes, nine were selected to perform a template picker. A more refined particle set was obtained using the previous templates, resulting in 808,253 particles. From this refined data set, seven classes consisting of a range of orientations of PAM_AP53_ were selected and a final template picker was run using these classes. The final particle set of PAM_AP53_- LVP bound to hPg contained 212,824 particles contained within 15 classes. Reconstruction was performed on the final particle set, followed by homogeneous refinement. Non-uniform refinement was performed to create a map focusing on increasing the resolution of lower secondary structural regions within PAM_AP53_. This resulted in a 4.4 Å map of PAM_AP53_-LVP bound to hPg. hPg from the finalized PAM_AP53_ structure was removed due to a lower resolution within this region making it incapable of being fully modeled. This is due to a number of PAM_AP53_ proteins not being bound to hPg. However, we clearly observed that there were no significant differences in structure in the 2D classes regardless of hPg being bound. The region of the map containing PAM_AP53_ was isolated and density modification of PAM_AP53_ was performed using Phenix refinement (16). This resulted in a 3.8 Å structure of PAM_AP53_ bound to hPg, a very good resolution considering the complexity of the particle.

The processes for which PAM_AP53_ was refined resulting in the final map is summarized in **Supplemental Figure 1S**. The 3.8 Å resolution was confirmed by the 0.143 cutoff FSC plot (**Supplemental Fig. 2S**). The data were submitted to EMDB and are available under the designation 41638. The full map containing the region of hPg bound to PAM protein is shown in **Supplemental Figure 3S**.

### Molecular modeling of PAM

The PAM_AP53_ cryo-EM map was used as the base to build the PAM_AP53_ model. The map was used to generate the initial model using the Phenix Map-to-Model function in order to obtain fragment placement in space. Using the ChimeraX (17) Build-Structure function, peptide fragments were generated based on the PAM_AP53_ sequence and placed according to the fragment placements generated from Phenix. Most peptide fragments of each domain of PAM_AP53_ were generated in ChimeraX with Φ: -57 and Ψ: -47 and flexed/flipped using ChimeraX ISOLDE tugging function. They were then connected using Phenix PDB editor and simulated to fit into the cryo-EM map using ChimeraX ISOLDE. The model was then further subjected to Phenix refinement and ChimeraX Trace-and-Build function. The process was repeated until a full model of the PAM_AP53_ structure was assembled.

The map (EMDB:41638) and finalized model of PAM_AP53_ (PDB:87VL) bound to hPg (**Fig. 4**) was obtained by maximally fitting PAM residues 19-356 into the cryo-EM map. Both the map (**Fig. 4A-B-C-D**) and the finalized model (**Fig. 4E-F-G-H**) are rotated 90° about a single axis to show different views of the overall tertiary structure of the protein and the individual domains that constitute the protein. The overall model shows a largely X-shaped structure. Residues 1-18 of PAM_AP53_ gave low electron density in the map and could not be fitted with a high level of confidence. The lower regions closer to the LVP membrane contained the Pro-Gly (P-G) and D- domains of PAM_AP53_ and the upper nodes contained the B- and C-domains and were present in the upper right region. The HVR and A-domain are located in the upper left region. PAM_AP53_ is modeled at a final length of 100 Å and a maximum and minimum width of 85 Å and 55 Å, respectively.

**Figure 4.**
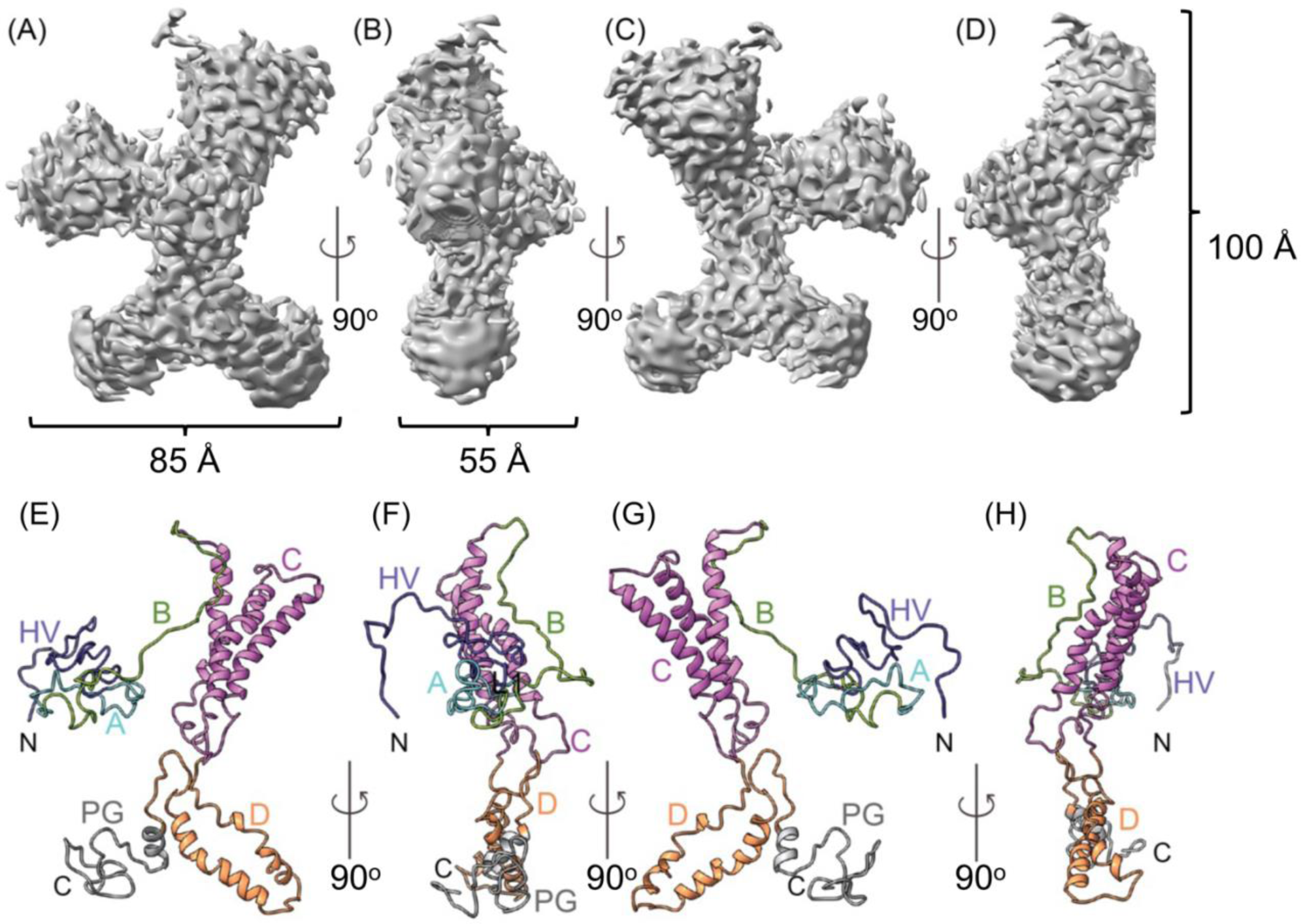
Cryo-EM map and model of PAM-hPg on the LVP. **(A-D).** The cryo-EM derived map of PAM_AP53_ resolved to 3.8 Å resolution, which was obtained from single particle analysis of PAM bound to hPg on the surface of LVPs. **(E-H).** PAM_AP53_ ribbon structure. The figure is depicting 90° rotation of the PAM map on the X-axis in each case. The PAM model showed an overall protein of 85 Å in width, 55 Å in thickness, and 100 Å in height. Each domain of PAM is colored differently; HVR in blue, A repeats (a1-a2) in cyan, B in green, C repeats (c1-c2-c3) in pink, D-domain in orange, and the Pro-Gly (PG) region in gray. The N- and C-termini for each model is denoted in black font.

### Overview of the 3D structure of PAM

The HVR and B-domains of PAM_AP53_ present as flexible regions with no defined secondary structural elements **(Fig. 4E-H).** The A-domain shows short regions of helical segments that are not well-defined due to the disruptive influence of the HVR and B-domains. The fitting of the C- and D-domains was prioritized due to clarity of the map. The C-domain is composed of three repeats, *viz.,* c1, c2, and c3, connected to each other by two loops of varying lengths **(Fig. 5A)**. Each c-repeat is folded into a right-handed α-helix and the repeats are further clustered together to form a three-helix bundle (THB) motif stabilized by hydrogen bonding and electrostatic interactions **(Fig. 5B)** as well as hydrophobic clustering **(Fig. 5C)**. The C-terminal portion of c3 and the N-terminal part of the D-domain exists as flexible loops connecting the C- and the D-domains (**Fig. 5D**). The latter, which forms a helix-loop-helix (HLH), is a helical domain (**Fig 5A,D**) that terminates in a flexible loop connecting it to the P-G-rich domain. This latter domain begins with a short helical segment. However, a larger portion of the domain exists as unstructured.

**Figure 5.**
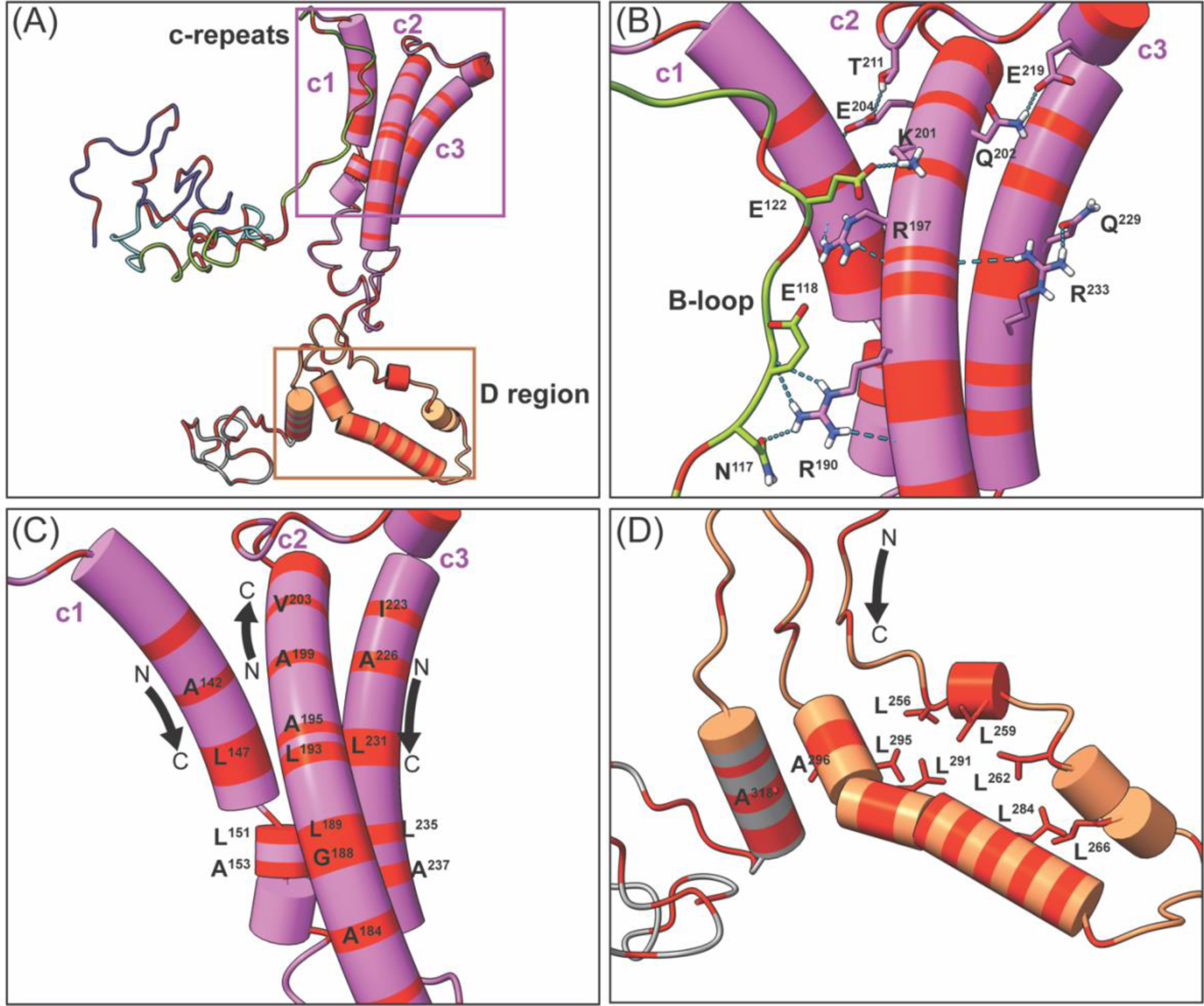
The α-helical regions of PAM_AP53_. Tube helices depicting the helix-helix interactions of the C-domain repeats and the D-domain. (**A).** The overall model of PAM highlighting the C-domain repeats (pink box) and D region (orange box). (**B).** Hydrogen bonds between helices involving residues R^197^, Q^202^, E^204^, T^211^ of c2 and E^219^, Q^229^, and R^233^ of c3. The positioning of the c2 helix was also stabilized by the B region loop involving residues K^201^ and R^190^ of c2 and N^117^, E^118^, and E^122^ of the B loop. **(C).** The c1-c2-c3 repeats are further stabilized by hydrophobic clustering of A^142^, L^147^, L^151^, A^153^ of c1, and A^184^, G^188^ L^189^, L^193^, A^195^, A^199^, and V^203^ of c2, and I^223^, A^226^, L^231^, L^235^, and A^237^ of c3. (**D).** The D-domain also shows helix-helix stabilization from clustering of L^256^, L^259^, L^262^, L^266^, L^284^, L^291^, A^295^, L^296^, and A^318^.

### Three helix bundle (THB) and helix-loop-helix (HLH) motifs of PAM

Significant portions of C- and D-domains of PAM consisted of heptad repeats (18). Heptad repeats favor helix formation and, in certain situations, hydrophobic residues of repeats of this type enable a single protein chain to fold back on itself (19) as seen in this case with PAM (**Fig 5C-D**). Here, the C-domain of PAM shows this property (**Fig. 5C**), largely stabilized by the clustering of Val and Leu residues occupying the positions **<u>a</u>** and **<u>d</u>** of the heptad repeat of PAM (18). Moreover, ∼70% of the Leu residues that stabilize the HLH of the D-domain (**Fig. 5D**) also occupy these same positions of the heptad repeat. Thus, the THB and HLH found in PAM_AP53_ are favored by its heptad repeats.

### Interactions between PAM C- and D-domains

The D-domain plays a large role in C-domain stabilization. As seen on the map and model of PAM_AP53_, the C-domain consists of tightly packed helix-helix interactions. The loop region between the c1-c2 helix as well as the region where c3 transitions into the D-region involves hydrophobic clustering of residues L^165^, A^166^, L^168^, A^170^, V^172^, V^175^, V^249^, L^249^, A^252^, L^302^, L ^305^, A^307^, and A^310^. Hydrogen bonding from residues T^169^, R^306^, K^309^, E^251^, S^254^, and N^253^ to the backbone of the C- and D-domains further stabilizes the interactions (**Fig. 6**).

**Figure 6.**
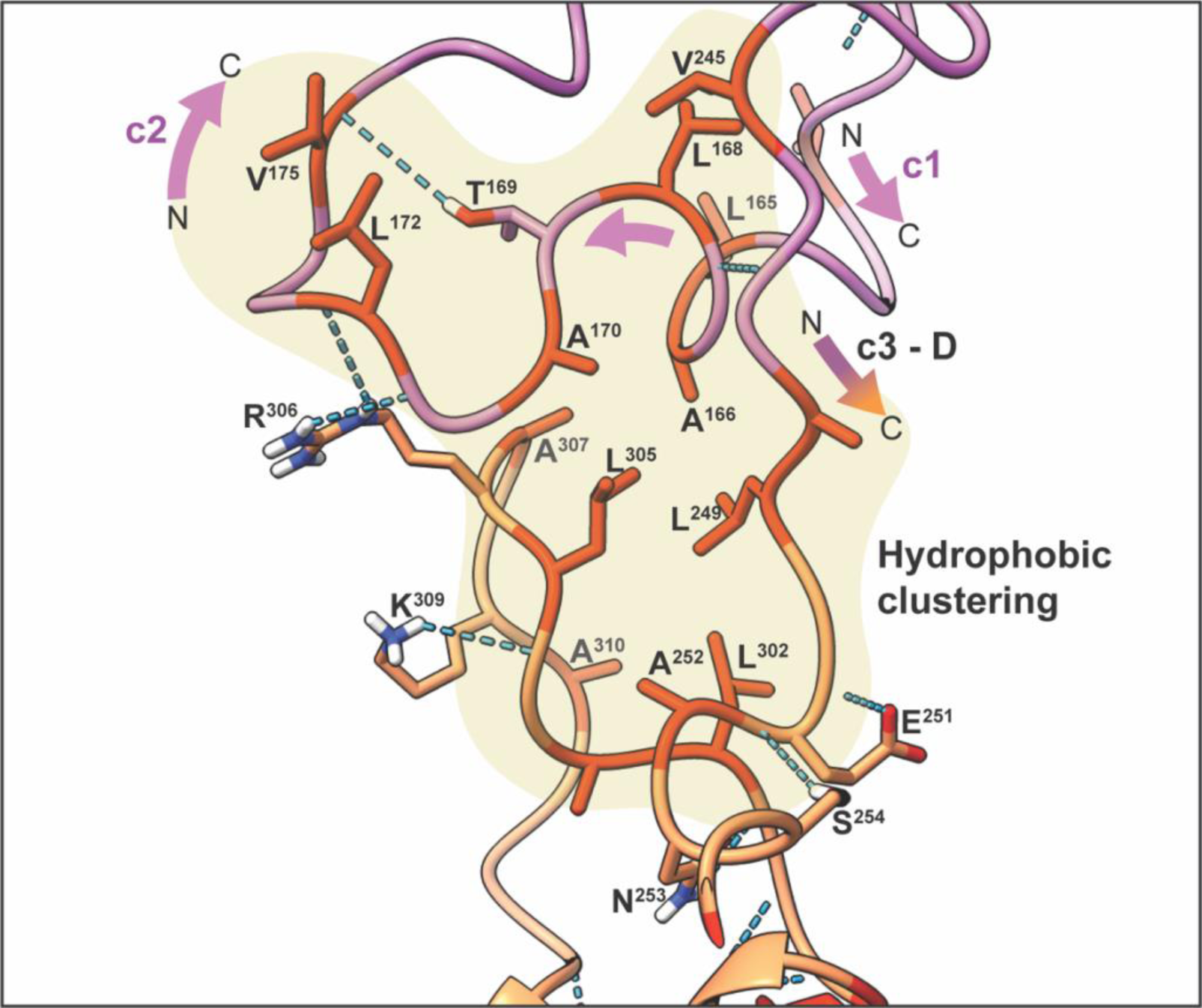
The interactions between C-domain repeats and the D-domain of PAM_AP53_. The backbone of the C-domain is shown in pink and that of the D-domain is depicted in orange, with hydrophobic residues shown in red for both regions. Hydrophobic clustering is highlighted in pale yellow between the C- and the D-domains. Stabilization of the C-D interactions involves residues hydrophobic clustering of L^165^, A^166^, L^168^, A^170^, V^172^, V^175^, V^249^, L^249^, A^252^, L^302^, L ^305^, A^307^, and A^310^, with hydrogen bonding from residues T^169^-V^175CO^, R^306^-L^172OC^, R^36^-E^171OC^, K^309^-K^304OC^, E^251^- E_251HN,_ S_254-_E_251OC,_ and N_253-_E_301OC._

### The A-domain of PAM_AP53_

The A-domain of PAM_AP53_ consists of two repeats indicated as a1 and a2 in **Figure 7A,B**. The Arg-His dipeptide motif, RH1 (R^72^H^73^E^74^E^75^) of a1, and the equivalent RH2 (R^85^H^86^D^87^H^88^D^89^) of the a2-repeat, have been previously shown to be critical residues in hPg-binding. This property is due to their side-chains forming a through-space isosteric lysine that interacts at the LBS of hPg- K2 (20). It is evident from **Figure 7** that the A-domain of PAM_AP53_ does not exist as an end-to-end α-helical structure, unlike the structures of truncated PAM_AP53_ peptides, such as VEK30, VEK50, and VEK75 (18, 20, 21) that are clearly shown to be full helices. **Figure 7C,D** highlights the intra A-domain and inter HVR, A-, and B-domain interactions that hinder α-helix formation in the A-domain in the intact protein. As shown, in the a1-repeat, A^63CO^ forms a hydrogen bond with R^67HN^ in a turn of an α-helix **(Fig. 7C**). However, the α-helix is disrupted and skewed by side-chain interactions between the HVR, A-, and B-domains, such as V^56CO^-K^69HN^, A^61OC^-R^67^, and R^67^- E^99OC^, leading to other intra A-domain interactions, such as those that exist between L^68OC^- E^71HN^ and N_70OC-_R_72HN._

**Figure 7.**
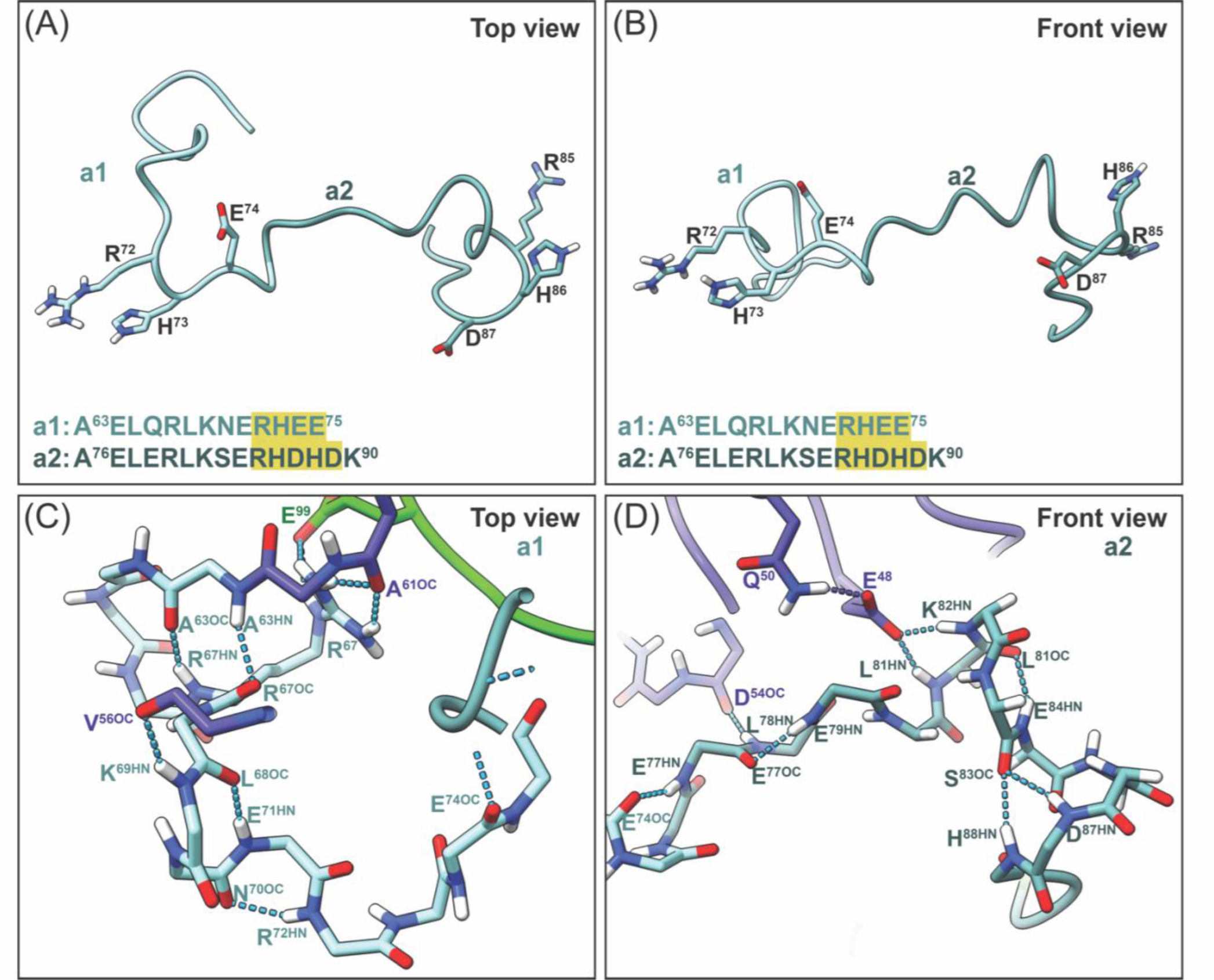
Interactions within the PAM_AP53_ a1a2 repeats. (**A-B).** Top view and front view of the a1 and a2 repeats of the A-domain highlighting the RH1 (light cyan) and RH2 (dark cyan) regions with isosteric lysine side-chain residues shown and highlighted in yellow. **(C).** The top view of the RH1 chain trace depicting the interaction between A^63HN^-R^67OC^, A^63OC^-R^67HN^, V^56OC^- K^69HN^, L^68OC^-E^71HN^, N^70O^-R^72H^, and residue R^67^-A^610C^-E^99^. **(D).** The front view of RH2 in chain trace depicting interactions between E^74OC^-E^77HN^, E^77OC^-E^79HN^, D^54OC^-L^78HN^, Q^50^-E^48^-K^82H^ and L_81HN,_ L_81OC-_E_84HN,_ and S_83OC-_D_78HN_ and H_88HN._

Similarly, within the a2-repeat, E^74^ of a1 begins a turn of an α-helix, but instead of hydrogen bonding with L^78OC^, it instead interacts with E^77HN^ of a2. This occurs due to the D^54OC^ of the HVR region interacting with L^78HN^ of the a2-repeat, as well as the E^48^ of the HVR interacting with L^81HN^ and K^82HN^ of the a1-repeat, making it possible for hydrogen bonding between E^77OC^-E^79HN^; L^81OC^- E^84NH^; S^83CO^-D^78HN^; and S^83CO^-H^88NH^ (**Fig. 7D**).

### Interactions between PAM and hPg-K2

The interaction between PAM and hPg-K2 was modeled using ChimeraX and ChimeraX- ISOLDE. The hPg-K2 (PDB:2DOH) was placed in space according to the full PAM-hPg cryo-EM map (**Fig. 8**), where PAM and hPg interact with each other. Using ISOLDE, hPg-K2 was simulated according to the map electron densities. The hydrogen bonding interactions can be seen between the side-chains of D^219^ and E^221^ of K2 with PAM side-chains of R^72^ and H^73^ (**Fig. 8**), in complete agreement with previous models using isolated hPg-K2 and the A-domain of PAM (20). Furthermore, appropriate distances were observed for interactions between the side-chain of R^234^ of hPg-K2 with the side-chain of E^74^ and with the backbone of K^69^ of PAM. The hydrogen bonding between N^70^ of the a1-repeat with E^162^ holds the c1 region in proximity, which allows electrostatic interactions between R^155^ of PAM and S^203^ of hPg-K2. Other electrostatic interactions can also be seen between Gln^50^ of the HVR of PAM and N^232^ of hPg-K2 (**Fig. 8**). The electrostatic interactions are based on the surface electrostatic map generated using the model in ChimeraX.

**Figure 8.**
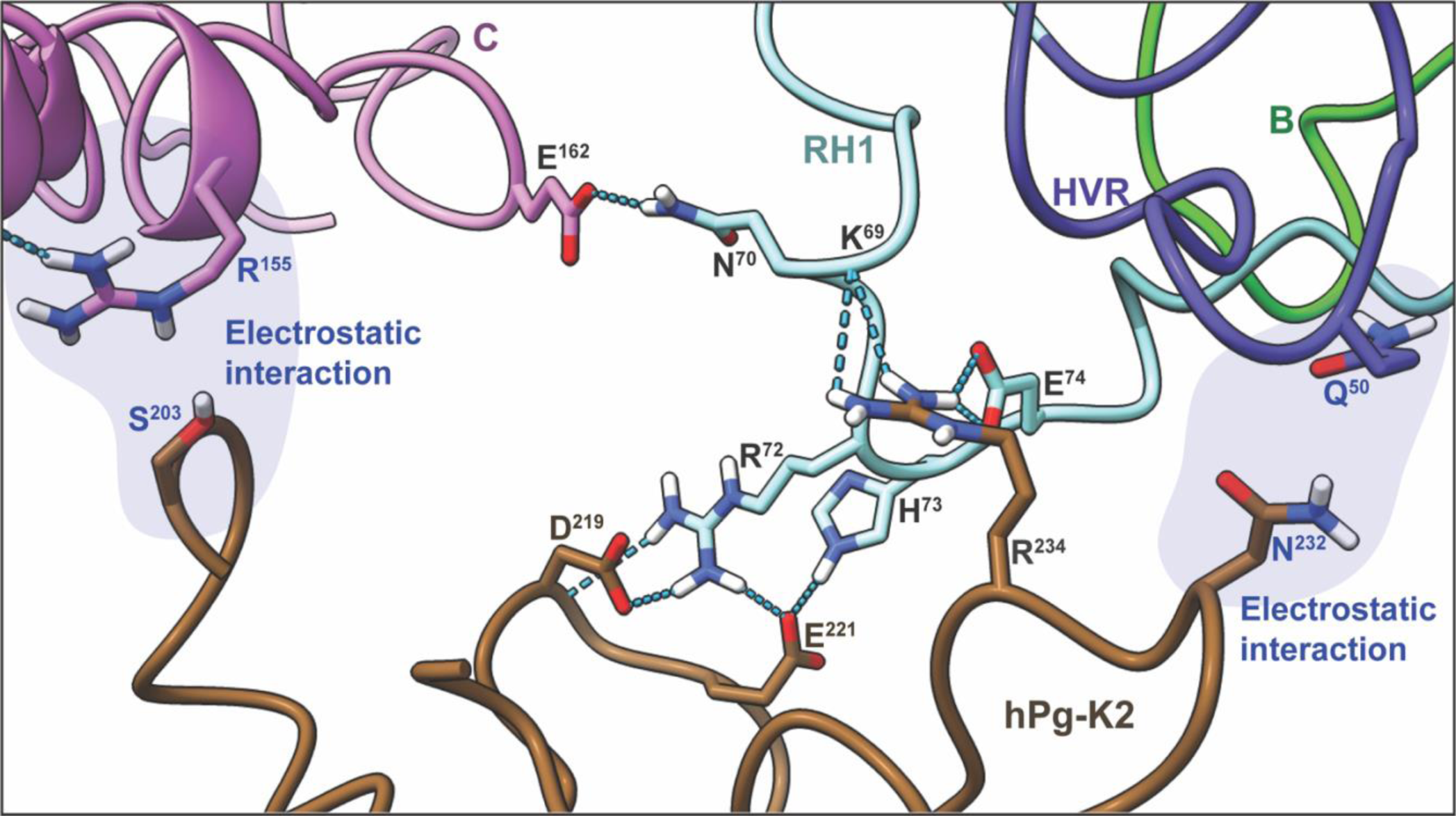
Interactions of hPg-K2 with PAM_AP53_-a1 in intact proteins. The placement of hPg- K2 from PDB:2DOH, based on the PAM/hPg cryo-EM map was simulated using ISOLDE to discover the interactions involved in complex formation. Side-chain hydrogen bonding was observed between D^219^ and E^221^ of hPg-K2 with R^72^ and H^73^ of PAM. Further backbone interactions between R^234^ of hPg-K2 and E^74^ and K^69^ of the PAM backbone were also observed. The side-chain residues of D^219^/E^221^ and R^234^ act as a lysine through-space isostere. Hydrogen bonding between N^70^ of RH1 with E^162^ holds the c1 region of the C-domain in proximity, allowing electrostatic interactions between R^155^ and S^203^ of hPg-K2. Other electrostatic interaction can also be seen between Q^50^ of PAM and N^232^ of K2.

## Discussion

Over 250 *emm-*encoded M-Prt types have been discovered, leading to the identification of an equivalent number of ∼250 GAS serotypes with distinct M-proteins (22, 23). Based on the number and arrangement of a short cluster of *emm* and *emm*-like genes, *viz., fcR*, *enn*, and *emm*, situated between the 5’-*mga* gene activator and 3’-*scpA* (collectively called the *mga* regulon), these M-Prt are organized into five Patterns, A-C, D, and E (24, 25). All direct hPg binding M-Prt, which we designate as PAMs, are present in Pattern D, which are skin-trophic strains. These different PAMs either have one or two a-repeats within the A-domain either of which can tightly bind to hPg in isolation. In the case of the PAM_AP53_, two possible hPg binding a-repeats, a1 and a2, are present. Aligned with this diversity in the *emm* gene, the primary structure of the gene product, especially in the HVR and A-C domains, exhibits a significant degree of sequence variation and functional differences as a GAS virulence factor. Nonetheless, M-Prts are organized in modular structural and functional domains, *viz.,* HVR, A, B, C, D, and P-G-rich regions, followed downstream by the LPST’G sortase A (srtA) cleavage motif, the transmembrane domain (TMD), and a short cytosolic (26). Major diversity exists among the GAS M-Prts within the HVR, A-, and B-domains, while the C-, D-, and P-G-domains, as well as the srtA cleavage region, the TMD, and the cytosolic peptide are much more conserved among M-Prts. The number of subdomains within these three domains (27) greatly influence the function of M-Prts, including GAS pathogenicity (28).

Our previous attempts at obtaining the cryo-EM structure of recombinant, soluble PAM proved to be too difficult due to its small size and poor contrast with the background. This led us to develop a novel method of incorporating recombinant proteins into LVPs with the goal of overcoming these complications. The addition of the VSV-G signal sequence and TMD+Cy C-terminal tail to the N- and C-terminal ends, respectively, in place of those regions of PAM, was sufficient to anchor the protein onto the surface of these LVPs while they were being produced. The LVP-containing PAM_AP53_ was assembled in growing human HEK-293 cells and the LVP isolated after integration of all components and amplification in these cells. This method could easily be applied to other membrane-bound proteins, bacterial or mammalian, whose structure is difficult to solve by cryo-EM.

### Tertiary structure of PAM

High-resolution structures of the amino-terminal fragments of M-Prts, including internal truncated portions of various M-Prts, with some as complexes to their binding partners, have been deposited in the PDB (12, 18, 20, 21, 29–32). However, to date, no known 3D membrane-bound structure of any full-length M-Prt exists. In order to more deeply probe the structure-function relationships of these types of M-Prts, we have solved, by cryo-EM, the 3D solution (vitreous) structure of the complete membrane-bound PAM_AP53_ structure. We focused on this particular M-Prt since it directly relates to our long-term interest in hPg and hPgRs. This particular receptor directly hijacks hPg from human host plasma, enables its rapid activation to hPm, and protects hPm from inactivation by natural inhibitors. These steps place a potent serine protease, hPm, on GAS cells, thus endowing these bacteria with the ability to invade host barriers to disseminate.

The novel strategy of integrating PAM on a LVP allows PAM_AP53_ to more closely mimic how M-Prts exist on a cell surface with the ectodomain projecting into solution. To our knowledge, this is the first bacterial surface-bound protein studied while anchored to a LVP membrane. Next, the anchored PAM_AP53_ was vitrified as a complex bound to hPg in order to enhance molecular weight compatibility with the cryo-EM technique. This also furthered enabled us to show the relevant portions of the interaction between membrane-bound PAM_AP53_ and hPg in intact proteins. While we could not resolve hPg to the same high degree as that of PAM_AP53_ (**Supplementary Fig. S3**), we nonetheless were able to map the known region of hPg (8, 11, 13, 33–35) that was bound to PAM by placing the 3D structure of the hPg-K2 domain within the confines of the experimentally-determined map coordinates.

As seen in our experimental model, the P-G region of PAM is a highly disordered structure due to the prevalence of Pro and Gly residues. The carboxyl-terminus of the P-G region, *viz.,* the TMD, is anchored to the membrane where these disordered structures then converge into a short helix wherein A^318^ clusters with A^296^ of the D-domain helix. A previous study of a truncated version of PAM, where residues 307-356 were missing, shows an increase in the helical content of the truncated PAM compared to the full-length PAM (18). This result further supports our model in that the HLH within the D-domain is destabilized by its interaction with the P-G region (**Fig. 5D**). Moreover, circular dichroism analysis of other shortened forms of PAM_AP53_, one containing only the HVR-A-B peptide, and another containing HVR-A-B-c1 peptide, shows that the former has higher helical content (18). This earlier finding, including the AlphaFold predicted structure of PAM_AP53_, suggests that the B-domain has the potential to be a full helix. Thus, consistent with the current 3D structure of PAM_AP53_, as reported for the CD spectrum HVR-A-B- c1 peptide, the interaction of the B-domain with the c1-repeat, represented in **Figure 5B**, eliminated helix formation in the B-domain.

As the P-G region proceeds upstream into the D-domain, the tail-end of the D-domain makes a hydrophobic cluster with a small loop separating the c1 and c2 repeats, as well as the junction connecting c3 to the D-region (**Fig. 6**). Overall, the P-G and the D-domains then stabilize the base of PAM_AP53_ with the C-domain loops, thus allowing the C-domain to fold into three helix bundles, bringing the D-domain and C-domain together as a short stump. These domain interactions between the B-C-D-domains and PG-region are major foundations of the compact tertiary structure of PAM_AP53_.

### Structural differences between M-Prts

Our cryo-EM structure of PAM_AP53_, (**Fig. 4** - the full map of hPg/PAM_AP53_ is shown in **Supplementary Fig. S3**) demonstrated that membrane-anchored PAM_AP53_ exists as a monomer with an extracellular projection of 100 Å. M-Prts are generally viewed as elongated tropomyosin-like proteins with no tertiary structural elements, even though only one such structure on a cell membrane has been observed by low resolution TEM (28). This latter M-Prt, M6_D471_ has major sequence variations in the virulence determinants of the HVR and A-C domains. From this TEM structure, M6_D471_ from a GAS Class A-C strain is believed to exist as a full α-helical protein, without discernable tertiary structure, forming coiled-coil dimers on the cell surface (28).

The main notable differences between M6_D471_ and PAM_AP53_ are their functions and their major human host binding partners. Unlike PAM_AP53_, M6_D471_ does not directly interact with hPg and, while it has a similar domain structure, considerable differences in sequence alignment are present. Amino acid sequence alignments of M6_D471_ and our PAM_AP53_ are shown in **Supplementary Figure S4**. According to these sequences, the most significant differences exist in the A-C-domains while strong homology exists in the D-, P-G, TM, and cytosolic domains of both proteins. It is notable that the A-domain of M6_D471_ does not contain amino acid residues that are critical for hPg binding, *viz.,* RH1 of the a1-repeat and RH2 of the a2-repeat, one or both of which is/are present in all Pattern D PAM proteins. M6_D471_ also contains a more extensive B-domain with more repeats than the PAM_AP53_ B-domain, wherein this region in PAM is largely disordered. The C-domain of M6_D471_ does not contain the c1-repeat which is critical for interaction between the C- and D-domains. Thus, the extension of this reasoning to M6_D471_ would result in a far longer and less ordered structure than is present in PAM_AP53_. The shortened extracellular projection obtained for PAM (100 Å), compared to the AlphaFold prediction of 440 Å for PAM_AP53_ and the TEM measurements of 500 Å for M6_D471_, can be explained by these sequence alignments in combination with the cryo-EM derived structure of PAM.

In the cryo-EM structure of PAM, the c1-repeat occupies a central position in favoring tertiary structure formation. The c1-repeat allows the formation of THB, which places c2 in the proper position to interact with the C-terminal loop of the short B-domain. Moreover, the interaction of the c1-c2 loop and the D-domain stabilizes the tertiary structure of PAM. The c1-repeat is not identifiable in M6_D471_, and, as such, the interactions stabilized by this repeat would be absent. Additionally, the residues seen to stabilize and promote the more compact structure of PAM are missing or mutated in M6_D471_ sequence (**Supplementary Fig. S4**). This would also destabilize any tertiary structure formation in M6. Further, the cryo-EM structure of PAM demonstrates that the B-domain is highly flexible and random.

Even though M6_D471_ is known to be fibrous, it is highly unlikely that it is a continuous end-to-end coil as depicted in the literature. Due to some conserved regions between PAM_AP53_ and M6_D471_, the most likely structure is a flexible P-G- and a coiled D-domain with more random elongated structures at the N-terminal HVR, A-, B-, and elongated helical C-domains resulting in the fibrous structure shown in the TEM micrographs. This would explain previous discrepancies in length of full-length M6, where it was found that M6_D471_ was ∼30% shorter experimentally than the predicted full-length M6_D471_ (36).

The M-Prt of all Pattern D GAS strains have evolved to bind directly to hPg as opposed to some pattern A-C strains, e.g., M6_D471_, which bind human fibrinogen (hFg). This functional difference appears to align with the structural difference between the experimentally obtained cryo-EM structure of Pattern D PAM and the Pattern A-C M6 structure. hFg forms a chain like hexameric structure that is ∼450 Å in length mirroring the structure of M6 **(Supplementary** Fig. 5). hPg is a more globular protein that is ∼100 Å and more closely resembles PAM_AP53_. Similarly, hFg forms a long coiled hexamer. This elongated structure of M6_D471_ is likely designed to maximize interactions with hFg. Consequently, it is plausible that differing M-Prts on the surface of GAS structurally evolved to allow the most efficient and stable binding of their human protein partners in order to employ the host as a partner in its pathogenicity.

### The A Domain

In our previous reductionist model of the PAM/hPg complex, the isolated A-domain exists as a helical peptide (11, 32). Herein we show that the fitting of the A-domain into the cryo-EM map, as well as the HVR and B-domain, unravels the helix. The overall helical content of 34% of the 3D structure is consistent with the CD spectrum of the intact PAM, which gives ∼30% helix in solution (18). This further strengthens the fact that the A-domain does not exist as a helix in intact PAM.

In the current 3D structure, the two RH motifs of the a-repeats are on the opposite sides of the polypeptide chain, similar to the reductionist models. Although both a-repeats are surface exposed and can bind to hPg with similar affinity in truncated PAM (32), our cryo-EM map shows that hPg preferentially interacts with the RH motif of the a1-repeat. This preferential binding to RH1 of a1 appears to be favored by the interaction between the a1 and the B-domain, which juxtaposes the lysine isosteric side-chains of a1 for hPg binding. Additionally, PAM is folded in such a way that N^70^ of a1 interacts with E^162^ of c1, bringing the a1-repeat closer to the C-domain. Therefore, the binding of hPg at the a1-repeat enables further stabilization of hPg through electrostatic interactions with the C-domain, including the HVR of PAM_AP53_. Moreover, with the proximity of a1 and the C-domain, other hPg domains, *viz.,* hPg-K3 and hPg-K5, are likely stabilized by the C- and D-domain **(Supplementary Fig. S3**). These stabilizing interactions are unlikely if hPg were bound at the a2-repeat.

In conclusion, our experimentally obtained cryo-EM map shows a novel compact structure of full-length membrane-bound PAM. Our structure sets a new paradigm on how the M-Prt folds. In the case that “form follows the function”, there are significant differences in some functions for different M-Prts, which seem to have evolved to maximize interactions with their major host binding partners. The compact structure of PAM is most likely similar in all Pattern D M-Prts, and these types of differences likely will be seen with other M-Prts given the high degree of sequence dissimilarities in the amino terminal regions of different GAS strains.

## Materials and Methods

### Construction of a PAM lentiviral particle (LVP) expression plasmid

A custom protein expression vector for mammalian cells was synthesized by Twist Bioscience (South San Francisco, CA) that contained the vesicular stomatitis virus glycoprotein (VSV-G) signal sequence (VSV-G_SS_; MKCLLYLAFLFIGVNC) followed by the VSV-G transmembrane domain + C-terminal cytosolic tail (VSV-G_TM+Cy_; FFFIIGLIIGLFLVLRVGIHLCIKLKHTKKRQIYTDIEMNRLGK). Two short linker (L) sequences encoding SGGGS were engineered to flank the internal ends of the VSV-G domains. The 5’-linker was followed by a 5’-NcoI (C’CATGG) site and the 3’-linker was preceded by a 3’-SacI (GAGCT’C) restriction site for cloning. The PAM protein coding sequence (residues 42- 397; 1-356 of the mature protein) was amplified by PCR from GAS_AP53_ genomic DNA using Q5 High-Fidelity DNA Polymerase (New England Biosciences, Ipswich, MA) with forward (5’- CCATGGAATAGAGCAGACGACGCTAGA-3’) and reverse (5’-GAGCTCTCACCTGTTGATGGTAACTGTCTC-3’) primers, respectively. This PCR product contained a 5’-NcoI site and 3’-SacI site for insertion into the expression plasmid. After accomplishing this, the final sequence (VSV-G_SS_-L-*NcoI*-PAM-*SacI*-L-VSV-G_TM+Cy_) was confirmed by Sanger sequencing. An endotoxin-free preparation of the cloned vector was made using the EndoFree Plasmid Maxi Kit (Qiagen, Hilden, DE).

### Expression and purification of PAM-LVP

For expression of PAM on the LVP surface, 8.5 µg of the expression plasmid, along with 5.5 µg each of HDM-Hgpm2 (NR-52517), HDM-tat1b (NR-52518), pRC-CMV-Rev1b (NR-52519), all obtained from BEI Resources in the SARS-Related Coronavirus 2, Wuhan-Hu-1 Spike-Pseudotyped Lentiviral Kit (NR-52948), were transfected into a 25 mL culture of Expi293 cells using the suggested protocol of the manufacturer. Three days post-transfection, PAM-LVP was harvested, diafiltered with PBS, and concentrated through a 500 kDa MWCO Amicon filter (Sigma-Aldrich, St. Louis, MO) under vacuum until a final volume of 20 µL was reached.

From the 20 µL concentrate, 5 µL was taken for a dot blot using an in-house polyclonal rabbit-anti PAM generated against the NH_2_-terminal domains of PAM. Free PAM was used as a positive control to obtain an estimate of the concentration of PAM on the LVP surface. Spike LVPs from the kit were used as a negative control to confirm that the antibody was specific for PAM. The remaining 15 μL of the PAM-LVP concentrate was frozen at -80° C until our scheduled time on the cryo-EM.

### Sample Preparation for Cryo-EM Analysis of PAM-LVP

The protocol for sample preparation of PAM-LVP was adapted from our previous published work (37) with the following changes. On the day of the cryo-EM experiments, the PAM-LVP concentrate was thawed at room temperature, after which recombinant hPg (3 µL of 0.6 mg/mL in 50 mM PBS, pH 7.4) was added to the 15 µL of concentrated PAM-LVP and the mixture was pipetted onto glow-discharged CF-1.2/1.3 Gold Ultrafoil® 300 mesh grids (Quantifoil, Jena, Germany). Five grids containing 3 μL each of the mixture were frozen and screened by TEM. The grid with the highest concentration of LVP was selected for the full cryo-EM run. The grids were blotted for 12 sec at a blot force setting of 10 at 25 °C/100% humidity. After this, the sample was plunge-frozen in liquid ethane for high-speed vitrification using the FEI Vitrobot MK IV (ThermoFisher). The grids containing the sample were then loaded into the FEI Titan Krios transmission electron microscope. Data were collected over 48 hr at 300 kV using the Gatan (Pleasanton, CA) K3 direct electron detector with a Gatan Quantum GIF energy filter in a super-resolution mode. Images were processed using EPU (ThermoFisher) software at a nominal magnification of 105,000X, resulting in a calibrated pixel size 0.437 Å. The total exposure dose was 55.00 e^-^/Å^2^, with a spherical aberration coefficient of 2.7 mm.

### Image processing and 3D mapping of cryo-EM images

The protocol for sample preparation of PAM-LVP was adapted from our previous publication (37) with the following changes. All image processing was performed on CryoSPARC v3.3.1 (http://www.cryosparc.com). The images were aligned and dose-weighted, and potential drift was removed by Patch Motion correction. The reference for focus correction was determined during the initial run. Contrast transfer function (CTF) was estimated for the dose-weighted micrographs using patch CTF estimation. Patch CTF extraction was performed to remove any micrographs that fell too far outside of the accepted focus or in which thick crystalline ice prevented data collection. The micrographs that did not contain LVPs were removed as they did not provide beneficial data.

### Resolution determination and structure validation

Map and model were validated using the Phenix Cryo-EM validation tool and confirmed by EMDB and PDB validation. The full finalized map and corresponding half-maps were analyzed using Fourier Shell Correlation (FSC) at a 0.143 threshold to determine the final resolution of the PAM-hPg map to be 3.8 Å. The fit between the cryo-EM map and the resulting model was also determined using FSC plotting at a 0.5 threshold giving a final fit of 3.6 Å between the two.

### Model building and refinement

Initial model building from the PAM cryo-EM map is similar to that previously described (37). Phenix was used to generate the initial model using map-to-model function with the known PAM sequence. Mature PAM protein lacked its signal peptide and residue numbering began from the HV region (N^1^) sequentially through the srtA LPST’G motif, immediately upstream of the TM domain of PAM. Segments of the PAM model were used as anchor points to place the remaining sequence using ChimeraX Peptide Build function. The placed peptides were then refined using ChimeraX ISOLDE (https://tristanic.github.io/isolde/) to match the placement of each residue and most importantly its backbone to the cryo-EM map. The fitted model was then further refined using Phenix trace-and-build function. The process was repeated multiple times and the final model then underwent final refinement using Phenix.

### Human plasminogen kringle-2 (hPg-K2)-placement, fitting, and simulation

hPg-K2 from PDB:2DOH was placed on the map in the region where PAM can be seen to be bound to hPg. The hPg-K2 model was then used for a top-down processing, fitting the interactions between hPg and PAM based on the cryo-EM map that was experimentally obtained. The fitting was accomplished through ChimeraX and the simulation of the model fitting into the electron density map was performed through ISOLDE. Part of the hPg-K2 backbone that was not covered very well in the density map was restrained manually in space.

