## Supplementary Data for "A high-resolution cryo-EM structure of a bacterial M-protein reveals a compact structure that diverges from related M-proteins"

### Supplementary Information

#### Supplementary Table S1. Cryo-EM Parameters

| Parameter | hPg-PAM-LVP<br>PDB: 8TVL, EMDB:41638 |
|---|---|
| Refinement program | Cryosparc |
| Magnification | 109,000 |
| Voltage, kV | 300 |
| Electron exposure, $e^-/Å^2$ | 55 |
| Defocus range | -1 to -3.2 |
| Pixel size, Å | 0.437 |
| Initial particle images | 808,253 |
| Final particle images | 212,824 |
| Symmetry imposed | C1 |
| Resolution unmasked,<br>FSC threshold 0.143 cutoff | 4.4 Å |
| **Model Design** | |
| Model resolution, Å | 3.6 |
| FSC threshold | 0.5 |
| Model resolution range | 6.9 |
| Map sharpening B-factor, $Å^2$ | -19 |
| Refinement program | Phenix, Chimera X |
| Number of atoms, non-H | 2,701 |
| Protein residues | 338 |
| Ligands | 0 |
| **B-factors, $Å^2$** | |
| Protein | 31 |
| **RMS deviations** | |
| Bond length, Å | 0.004 |
| Bond angle, $^o$ | 0.866 |
| **Validation** | |
| Mol probity score | 2.53 |
| Clash score | 14.96 |
| Poor rotamers, % | 0 |
| Ramachandran favored, % | 73.81 |
| Ramachandran allowed, & | 25.30 |
| Ramachandran disallowed, % | 0.89 |
| EMringer score | 2.49 |

**Supplementary Figure S1**

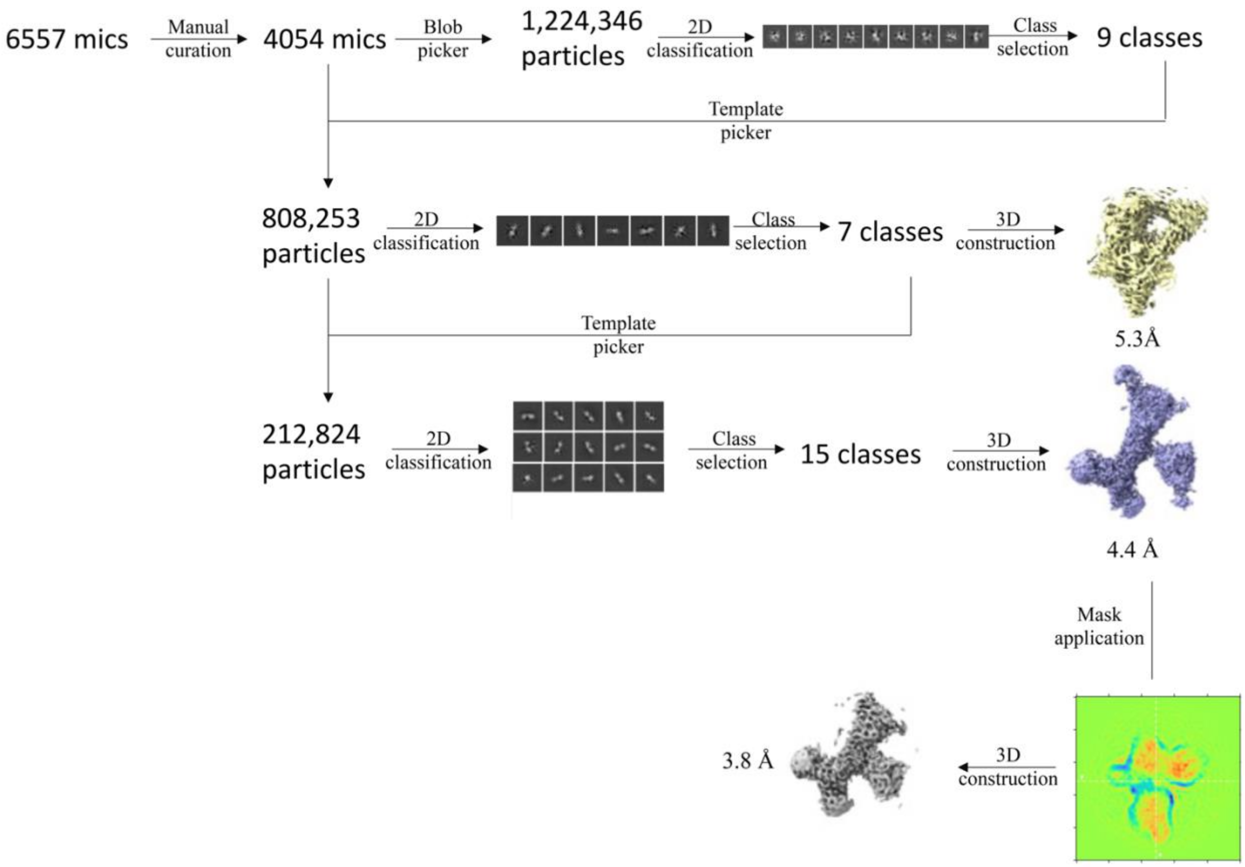


**Supplementary Figure S1.** Workflow diagram describing the procedure for generating the finalized map structure of PAM bound to hPg in Cryosparc. The resolution of the maps was determined at FSC 0.143.

**Supplementary Figure S2**

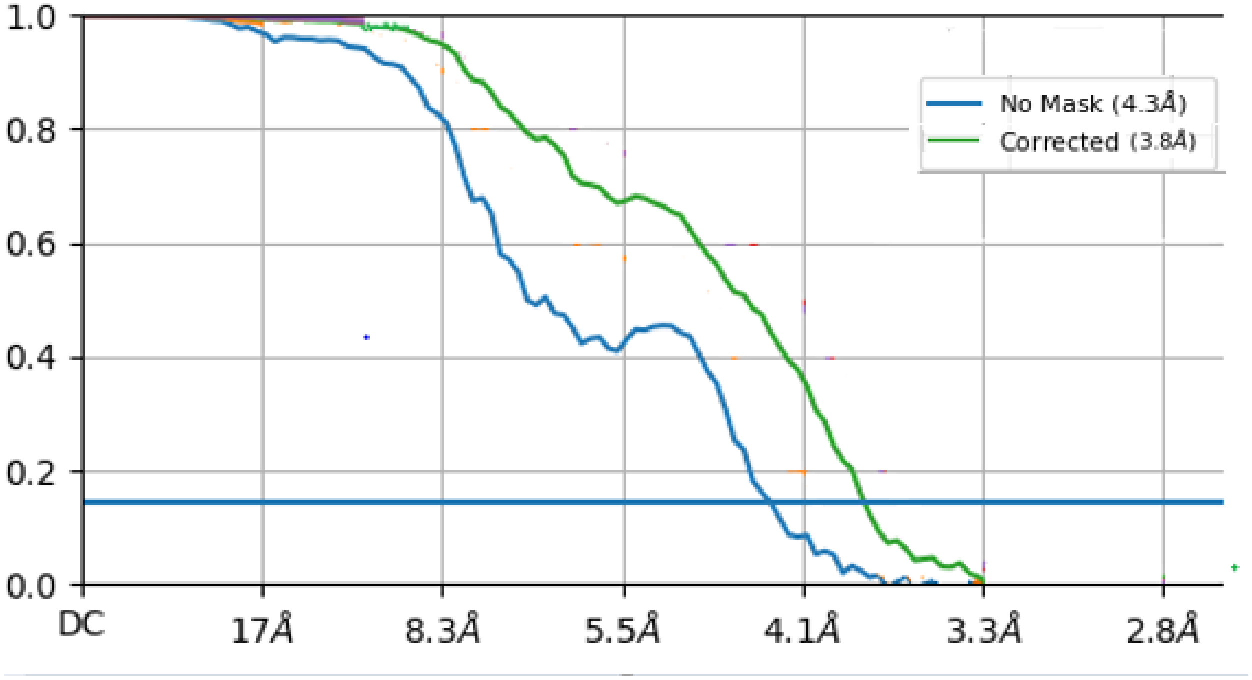


**Supplementary Figure S2.** Supplementary Figure S2. FSC plot of the finalized $PAM_{AP53}$ map. Values shown contain both the unasked and final fitted resolutions for the PAM map using the gold standard FSC value of 0.143.

**Supplementary Figure S3**

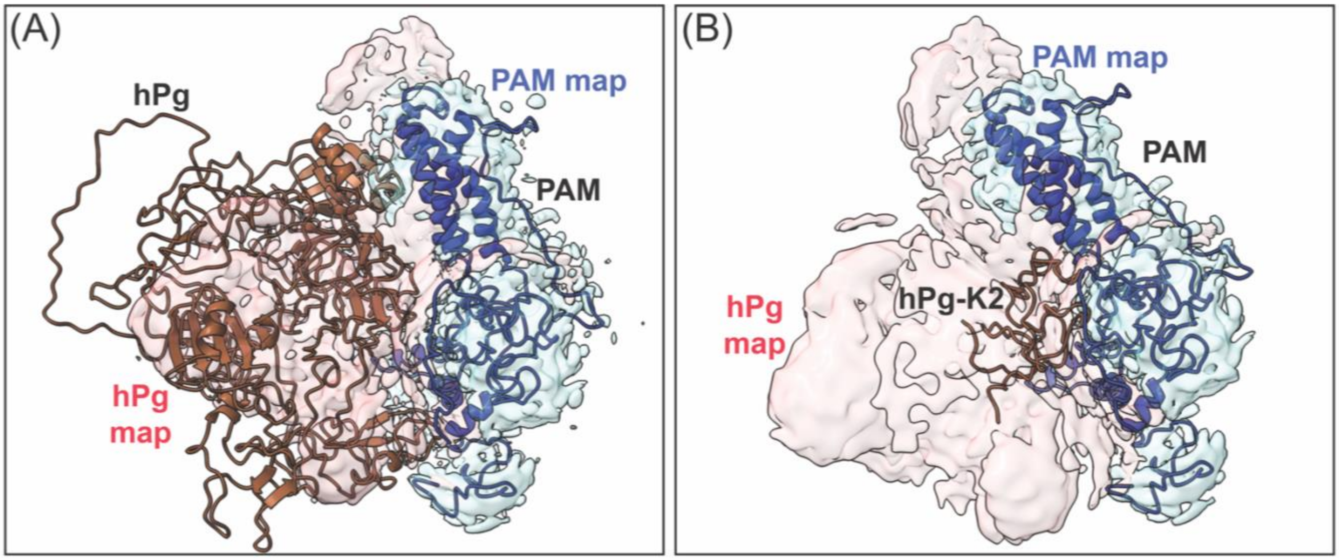


**Supplementary Figure S3. Full map of the PAM-hPg complex.** This map was used to fit the K2 region to the a1 binding site within PAM. **(A)** The rough placement of closed hPg (brown) against PAM (blue) model into the hPg map region of PAM-hPg complex. **(B)** The fitted and simulated hPg-K2 into the PAM-hPg cryo-EM map. Due to variability of PAM not being 100% bound to hPg and the highly flexible nature of the open conformation of hPg this region could not be fully resolved. Overall, the PAM structure was not shown to vary to a significant degree whether or not hPg was bound.

**Supplementary Figure S4**

```
emm6        RVFPRGTVENPDKARELLNKYDVENSMLQANNDKLTTENNNLTDQNKNLTTENKNLTDQN     60
emm53       --------NRADDA---------RNEVLRGN----------LV-----------------     16
                    :. *.*         .*.:*:.*          *.

emm6        KNLTTENKNLTDQNKNLTTENKELKAEENRLTTENKGLTKKLSEAEEEAANKERENKE--     118
emm53       ----RAEL------------WYRQIQENDQLKLENKGLKTDLREKEEELQGLKDDVEKLT     60
                  :               .   *:::*. *****...* * ***  . : : ::

emm6        ---AIGTLKKTLD-ETVKDKIAKEQESKETIGTLKKTLDETVKDKIAKEQESKETIGTLK     174
emm53       ADAELQRLKNERHEEAELERLKSERHDHDKKEAERKAL----EDKLADKQEH--LNGALR     114
                :  **:  . *:  ::: .*:..::.  : :*:*    :**:*.:**     *:*:

emm6        KTLDETVKDKIAKEQ----------ESKETIGTLKKILDETVKDKIAREQKSKQDIGALK     224
emm53       -YINEKEAERKEKEAEQKKLKEEKQISDASRQGLRRDLDA----SREAKKQVEKDLANLT     169
              ::*.  ::  **            *. :   *:: **     .   ::: ::*:. *.

emm6        QELAKKDEGNKVSEASRKGLRRDLDASREAKKQVEKDLANLTAELDKVKEEKQISDASRQ     284
emm53       AELDKVKEEKQISDASRQGLRRDLDASREAKKQVEKGLANLTAELDKVKEEKQISDASRQ     229
             ** * .* :::*:***:******************.***********************

emm6        GLRRDLDASREAKKQVEKALEEANSKLAALEKLNKELEESKKLTEKEKAELQAKLEAEAK     344
emm53       GLRRDLDASREAKKQVEKALEEANSKLAALEKLNKELEESKKLTEKEKAELQAKLEAEAK     289
            ************************************************************

emm6        ALKEQLAKQAEELAKLRAGKASDSQTPDAKPGNKVVPGKGQAPQAGTKPNQNKAPMKETK     404
emm53       ALKEQLAKQAEELAKLRAEKASDSQTPDAKPGNKAVPGKGQAPQAGTKPNQNKAPMKETK     349
            ****************** ***************.*************************

emm6        RQLPSTGETANPFFTAAALTVMATAGVAAVVKRKEEN   441
emm53       RQLPSTGEAANPFFTAAALTVMATAGVAAVVKRKEEN   386
            ********:****************************
```

**Supplementary Figure S4. The alignment of amino acid sequences of M6D471 *(emm6)* with PAMAP53 *(emm53)*.** Each domain of PAM is colored similar to those of **Figure 4**, dark blue for HVR, cyan for the A-domain, green for the B-domain, pink for the C-domain, orange for the D-domain, and gray for the Pro-Gly region. The sequence of the c3 repeat, the D-domain and Pro-Gly region is highly conserved. The hydrophobic residues responsible for C- and D-domain folding by clustering are highlighted in yellow, the residues interaction between B- and C- domain in green, and residues responsible for bringing the A-domain in juxtaposition to the C-domain are highlighted in cyan.

**Supplementary Figure S5**

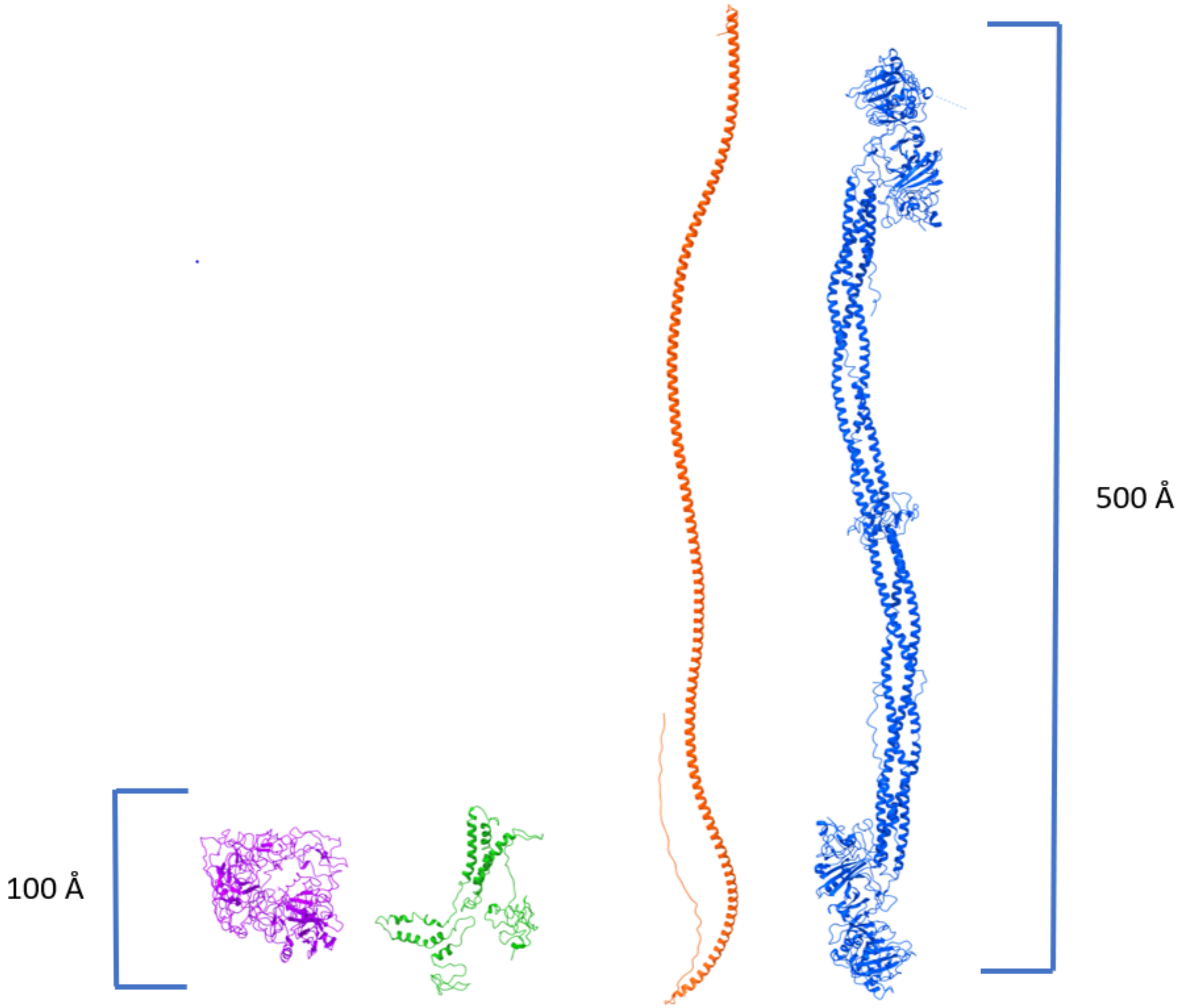


**Supplemental Figure S5. The M-Prt within GAS mirrors its binding partner.** The AlphaFold structure of $M6_{D471}$ (AF-P08089; red) is a largely linearized protein roughly 500 Å long protein matching its binding partner fibrinogen (PDB:3GHG; blue). $PAM_{AP53}$ (PDB:87VL, green) contrastingly is a more compact protein ~100 nm in length to maximize potential interactions with the globular hPg (PDB:4DUU; purple).
